# Decoupling of rates of protein synthesis from cell expansion leads to supergrowth

**DOI:** 10.1101/498600

**Authors:** Benjamin D. Knapp, Pascal Odermatt, Enrique R. Rojas, Wenpeng Cheng, Xiangwei He, Kerwyn Casey Huang, Fred Chang

## Abstract

Cell growth is a complex process in which cells synthesize cellular components while they increase in size. It is generally assumed that the rate of biosynthesis must somehow be coordinated with the rate of growth in order to maintain intracellular concentrations. However, little is known about potential feedback mechanisms that could achieve proteome homeostasis, or the consequences when this homeostasis is perturbed. Here, we identified conditions in which fission yeast cells are prevented from volume expansion but nevertheless continue to synthesize biomass, leading to global accumulation of proteins and increased cytoplasmic density. Upon removal of these perturbations, this biomass accumulation drove cells to undergo a multi-generational period of “supergrowth” in which rapid volume growth outpaced biosynthesis, returning proteome concentrations back to normal within hours. These findings demonstrate a novel mechanism for global proteome homeostasis based on modulation of volume growth and dilution.

## INTRODUCTION

Proliferating cells generally increase their biomass and volume during the cell cycle before dividing. Although much is understood about how duplication of certain cellular components such as the chromosomes is accomplished, much less is known about how the proteome itself is duplicated. The concentrations of many proteins are thought to be maintained during cell growth^1–3^. While global mechanisms of proteome homeostasis have been identified^4^, it is unknown the extent to which the concentrations of individual proteins are coordinated with cell volume^5^, whether this coordination can be altered, and the effects of such a perturbation.

The rate of cellular growth, which we define here as the increase of cellular volume, is determined by numerous factors. The biosynthesis of cellular components has been speculated to set growth rate, as decreasing protein translation, for instance, can slow or halt growth. The rate of cellular growth is also affected by cell size; many organisms, including bacteria, fungi, and mammalian cells, exhibit exponential growth^6,7^ in which the absolute growth rate at steady-state is proportional to cell size. Although the mechanism(s) for achieving exponential growth remains to be determined, it may arise from the scaling of the biosynthetic machinery with cell size: if protein synthesis is coupled to cell size, and cell size is dictated by protein concentration, then exponential growth will result.

The fission yeast *Schizosaccharomyces pombe* is an established model for cell-cycle regulation and growth. The simple, rod-shaped morphology and regular growth and division patterns of these cells make them highly amenable to quantitative studies. During their cell cycle, *S. pombe* cells exhibit polarized tip growth at one or both cell tips during interphase^8,9^, and growth halts during mitosis and cytokinesis^10,11^. Like other tip-growing cells, the growth of the cell surface is directed by polarity machinery that ultimately mediates remodeling and insertion of new cell wall at the cell tips. Growth of the surface is further impacted by mechanical factors, such as the turgor pressure due to osmolyte concentration imbalances across the membrane that expands the elastic cell wall^8,10^. While exponential volume growth at the single-cell level has been observed in many cell types, whether individual fission yeast cells exhibit such behavior has been a source of controversy for decades; the current consensus is a bilinear growth behavior with an increased slope later in the cell cycle^11–16^.

Biomass synthesis (largely driven by protein synthesis) and volume increase (driven by membrane and cell-wall synthesis) must be coordinated during growth, but it is unknown how this fundamental coupling is achieved. Here, we present multiple ways of uncoupling the coordination between biomass and volume in *S. pombe* cells, and thus provide new insights into the role of proteome homeostasis in cell growth. We first establish that fission yeast cells exhibit cell size-dependent, exponential growth for a large fraction of the cell cycle. Through manipulation of turgor pressure or secretion, we decouple the rate of global protein synthesis from growth rate. These conditions produce a global excess of proteins within the cytoplasm, which in turn results in an extended period of extremely rapid growth. These findings thus provide new insights into the role of cytoplasmic density in cell growth and proteome homeostasis, whereby the rapid expansion of dense cells accelerates the re-equilibration of protein concentrations.

## RESULTS

### Fission yeast cells grow exponentially during a large fraction of the cell cycle

To determine the growth behavior of individual *S. pombe* cells, we imaged cells in time lapse in microfluidic chambers and implemented automated image analysis methods to quantify cellular dimensions at subpixel resolution (Figure 1A, S1A; Methods). In agreement with previous studies, these rod-shaped cells elongated from ~8 μm to 14 μm before entering mitosis (Figure 1B, S1B), and maintained a constant width of ~4 μm (Figure 1B, S1C). Instantaneous growth rates (defined as *dL/dt*, where *L* is cell length) of wild-type cells ranged from 2−5 μm/h at 30°C (Figure 1C), with cell length highly correlated with elongation rate, indicating that growth rate accelerates during the cell cycle. In purely exponential growth, absolute growth rate scales linearly with cell size, hence 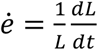 (size-normalized growth rate) is a constant. Quantification of *ė* revealed that fission yeast growth during the cell cycle is organized into at least three phases (Figure 1D): (I) growth acceleration for the first ~10−20% of the cell cycle (immediately after cell separation), (II) exponential growth for the majority of interphase, in which *ė* = 0.35 ± 0.06 h^−1^ (corresponding to doubling biomass during interphase in 2.0±0.3 h), and (III) deceleration to zero growth in mitosis and cytokinesis. These data are inconsistent with simple linear or bilinear growth models (Figure S2). We further tested the effect of cell size on growth rate by analyzing abnormally large cells (*cdc25-22* cells at the semi-permissive temperature 30°C, which divide at 30±4 μm in length). These large cells grew much faster than wild-type cells and followed a similar scaling of single-cell elongation rate with size, with a size-normalized growth rate *ė* of 30 ± 0.05 h^−1^ (Figure 1C, S1D,E). These findings establish exponential growth behavior in fission yeast, with absolute growth rates that scale with cell size over a wide range of volumes.

**Figure 1:**
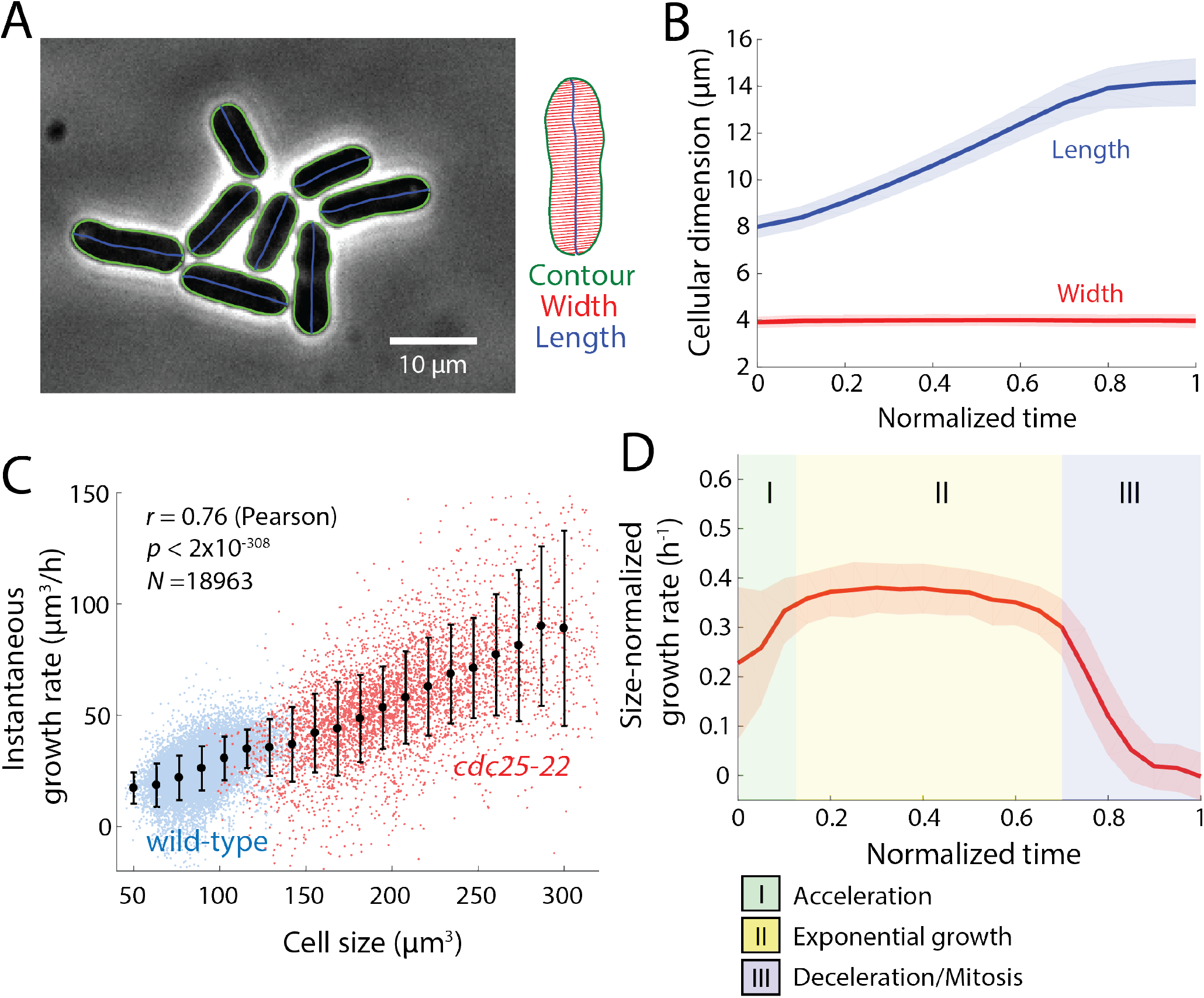
*S. pombe* cells grow exponentially. A. Wild-type (WT) fission yeast cells imaged in phase contrast. Green outlines were obtained via automated segmentation (Methods). Manual inspection revealed that segmentation was accurate for virtually all cells, regardless of cell crowding.
B. Population-averaged length and width of growing fission yeast cells, measured as a function of time after the mechanical separation of daughter cells after septation normalized by cell-cycle time (n = 1129 cells). Solid lines and shaded areas represent the mean ± 1 standard deviation (S.D.). Cell width was essentially invariant, while length increased over the cell cycle until the cell reaches the division length of ~14 μm.
C. Instantaneous growth rate *dL/dt* increased with cell length. The trend for wild-type (WT) cells (blue) continued for *cdc25-22* cells (red), which are substantially longer. Black lines represent best linear fits for each strain (wildtype, n=1129 cells; *cdc25-22, n*=246 cells).
D. Instantaneous size-normalized growth rate (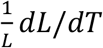) of wild-type cells, as a function of normalized cell-cycle time as in (B). The size-normalized growth rate was constant for a large fraction of the cell cycle, signifying growth proportional to size (exponential growth). Cells had three phases of growth: (I) super-exponential acceleration, (II) steady-state exponential growth, and (III) deceleration as cells approach mitosis.

### Cells grow slowly during oscillatory osmotic shocks, and exhibit super-fast growth upon exit from oscillations

In terms of the mechanics of cell growth, internal turgor pressure provides physical force for expansion of the cell wall to increase cellular volume. Fission yeast cells have a thick (~200 nm), elastic cell wall inflated by high turgor pressures of ~1.5 MPa (15 atm)^10,17^. Examining the effects of adding the osmotic agent D-sorbitol (hereafter sorbitol) to the media has proven useful for probing the mechanical effects of reducing turgor pressure^10,18–20^. We investigated the effect of mechanical perturbations on cell growth using osmotic oscillations in repeated cycles of hyper- and hypoosmotic shocks (Methods)^21^. We exposed *S. pombe* cells in a microfluidic device to repeated switches between rich medium (YE5S) and YE5S+0.5 M D-sorbitol (Figure 2A, S3A). In these initial experiments, we applied 24 oscillatory cycles of 0.5 M sorbitol shocks with a 10-min period for 4 hours, then subsequently followed cells in YE5S media without sorbitol. Each acute addition of 0.5 M sorbitol caused rapid water efflux, loss of turgor pressure, and shrinkage in volume by ~20%, with mean longitudinal and radial contractions in cell size of 3.5% and 7%, respectively (Figure 2B)^10^. During each 5-min hyperosmotic period, cells partially re-inflated, dependent on osmotic adaption mechanisms (Figure 2B, S3B). Upon hypoosmotic shifts back to YE5S medium without sorbitol, cells rapidly returned to their normal width in the absence of shocks (Figure 2B, S3B), suggesting that *S. pombe* cells can rapidly downregulate turgor pressure to a preferred value when envelope stresses increase above a certain point.

**Figure 2:**
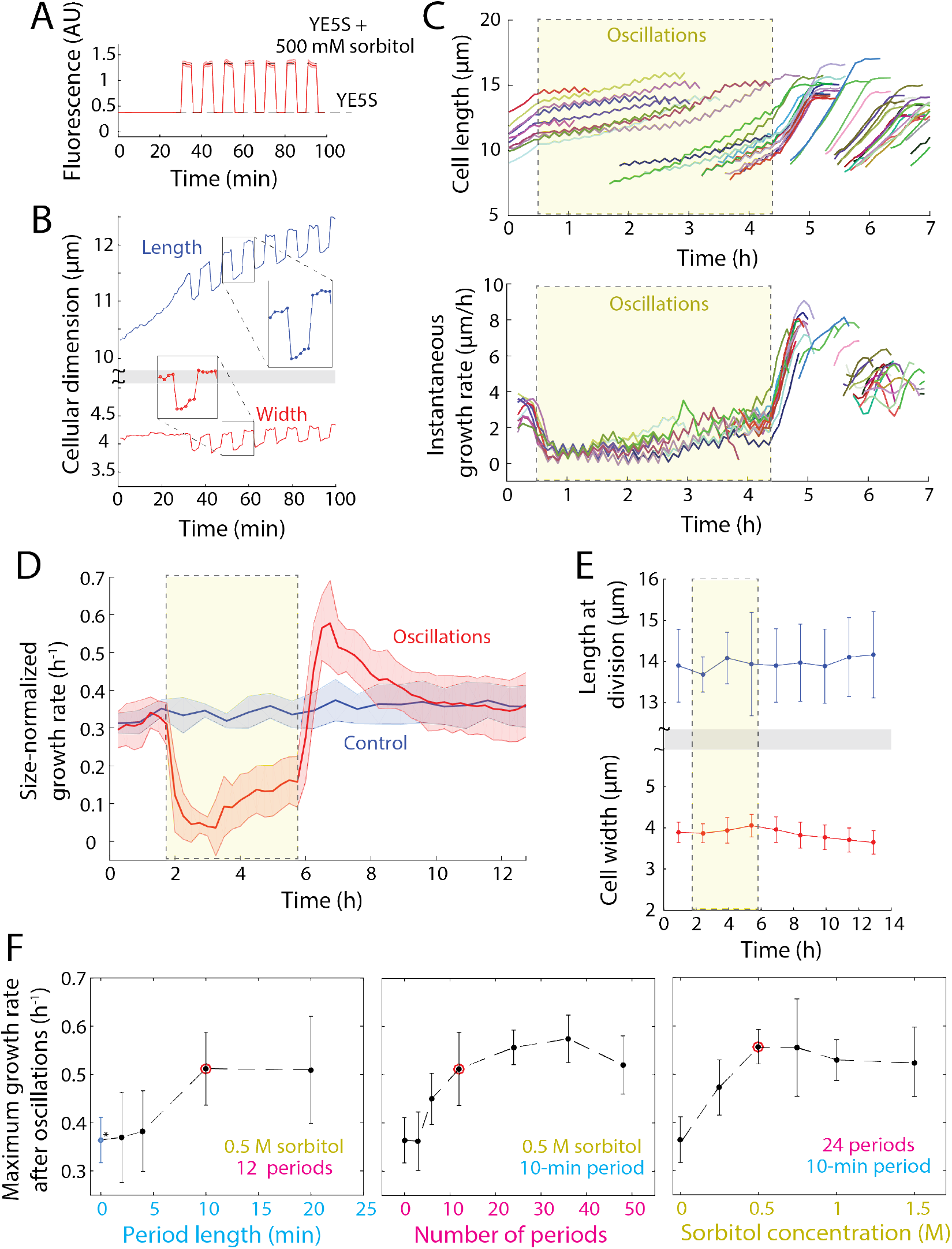
Exit from osmotic shock oscillations induces robust supergrowth. A. Cells were equilibrated to achieve steady-state growth in YE5S, and then exposed to cycles of hyperosmotic and hypoosmotic shocks with 500 mM sorbitol, with 5 min between shocks.
B. Length of a representative wild-type (WT) cell increased more rapidly during periods in YE5S+500 mM sorbitol, but returned to the value expected of growth in YE5S after hypoosmotic shocks. Width increased in YE5S+500 mM sorbitol, and was maintained at an approximately constant level in YE5S.
C. Length (top) and instantaneous growth rates (bottom) of wild-type cells (n = 45) before, during (yellow shading), and after 24 cycles of 0.5-M sorbitol osmotic shock oscillations with 10-min period. Cells are the same color in both plots. Cells were imaged every 5 min. After the oscillations, cells exhibited faster growth (supergrowth) for multiple generations. Traces that begin at later times points are newly formed daughter cells.
D. Size growth rates of wild-type cells throughout an oscillatory osmotic shock experiment (red, *n* = 973 cells). Growth rate during oscillations (yellow, dashed box) was lower than the control that was kept in YE5S throughout (blue, *n* = 1242). Growth rate increased after cells exited the oscillations, decreasing back to the control growth rate with a time constant of 1.3 h.
E. Cell-size control was unaffected by oscillations or supergrowth, with constant length (top) and width (bottom) at the time of division throughout the experiment in (C).
F. Maximal size-normalized growth rates during supergrowth as a function of (i) oscillatory shock period length, (ii) number of periods, and (iii) sorbitol concentration (n = 46−536 cells per data point). Each parameter was varied while the other two parameters were held fixed. Red circles represent our standard conditions of 24 cycles of 500-mM shocks with 10-min period. Error bars represent one standard deviation.

Over the course of 4 hours of these osmotic cycles, cells remained viable and continued to grow and divide (Movie S1, Figure S3C). However, the mean growth rate was lower than that of control cells: growth initially slowed (Figure 2C) for the first ~2 h (to <0.06 h^−1^, Figure 2D), followed by adaptation to the oscillations after ~3 h that resulted in a size-normalized growth rate increase to ~0.2 h^−1^ (Figure 2D). When cells were returned after the cycles of osmotic oscillations to YE5S without any sorbitol, they grew unusually fast, with tip growth rates >8 μm/h (population mean 6.5±0.9 μm/h), almost two-fold higher than mean control tip growth rates (3.8 μm/hr, Figure 2C); we refer to this period of unprecedented rapid growth as “supergrowth.” Size-normalized growth rates indicated that cells were growing abnormally rapidly for their size (*ė* = 0.6 h^−1^ vs. 0.35 h^−1^ for control cells, Figure 2D). These growth behaviors were highly stereotypical throughout the population (>96% cells, 49/51), and were substantially faster than any previously reported growth rates for fission yeast. Acceleration began by 15 minutes after exit from oscillations (Figure 2C), and growth rate reached a peak just before cells entered mitosis (Figure 2C, S4A). Elevated growth rates persisted for 2-3 generations (~4 hours, Figure 2C,D, S4B), but in each cell cycle, the rate decreased in a step-wise manner from the previous generation even though cells still exhibited exponential growth (Figure S4B,C). Growth still decelerated normally during mitosis and cytokinesis. Interestingly, the initial acceleration phase of the cell cycle (Figure 1D) was largely absent during supergrowth (Figure S4B), suggesting that cells were primed for exponential growth immediately after division.

Despite the large changes in growth rate during and after the oscillations, cell morphology and size were remarkably normal. Cells exhibited tip growth during interphase without large changes in cell width or tip shape (Figure S5). Importantly, cells entered mitosis at the normal cell length of 14 μm during oscillations and during supergrowth despite a >10-fold range in growth rates, indicating that changes in growth rate did not affect cell-size control (Figure 2E). During supergrowth, cell-cycle periods were shorter because due to a decrease in the duration of interphase, but the durations of mitosis and cytokinesis were normal, indicating that not all cellular processes were sped up (Figure S4C,D). These data suggest that cell size and the periods of cell division are controlled independently of growth rate.

To determine the requirements for supergrowth, we examined its dependence on oscillation parameters (amplitude, period, number of periods). We systematically varied each parameter around our baseline values (0.5 M, 10-min period, 24 periods), keeping the other two parameters fixed. In general, the maximal supergrowth rate increased gradually with increasing amplitude, period, or number of periods (Figure 2F). Single shocks were generally not sufficient (Figure 2Fii). Oscillations with 1 M sorbitol, which caused ~50% volume loss, resulted in a near-cessation of growth during oscillations, but then subsequently led to rapid supergrowth (Figure 2Fiii). These data suggest that growth and cell-cycle progression during oscillations are not required for supergrowth. The graded effects suggest that the rapid growth state is not regulated by an all-or-nothing switch, but rather accumulates over time during the osmotic shifts.

### Supergrowth is independent of established osmotic stress and growth pathways

To probe the mechanism underlying supergrowth, we considered two non-exclusive models. The first model involves the osmotic stress-response pathway, which could signal to growth pathways to regulate growth rate. The second model is based on material storage, in which components important for growth accumulate during periods of slow volume expansion because biosynthesis does not slow commensurately with growth rate. The excess of these materials upon exit from oscillations could then drive the subsequent supergrowth.

As an initial test of the signaling model, we assayed various mutants for their ability to undergo supergrowth. We assessed strains lacking Sty1 and Pmk1 (MAP kinases at the hub of the pathways that response to osmotic, cell-wall, and other environmental stresses)^22,23^; Gpd1, which regulates glycerol synthesis during turgor adaptation to osmotic stresses^19,22^; and Cch1, a calcium channel involved in response to cell-wall stresses^24^. We also queried regulators in the TOR pathway, a central regulator of growth: Ssp2 (an AMPK-like protein kinase that regulates TORC1 activity)^25,26^ and Gad8 (an AGC protein kinase effector of TORC2)^27^. All these mutants exhibited supergrowth responses similar to that of wildtype after oscillatory osmotic shock (Figure S6). Ribosomal mutants with slower basal growth rates showed a proportional supergrowth response (Figure S6). Thus, these signaling pathways regulating cellular responses to stress, turgor, and growth are not required for supergrowth.

### The proteome globally increases in concentration during osmotic oscillations

The material storage model predicts that components important for growth rise in concentration during oscillations. As an initial test, we monitored a fluorescent protein marker, E2-mCrimson expressed from the *ACT1* (actin gene) promoter. E2-mCrimson is stable and folds relatively rapidly^28^. Under normal growth, mCrimson intensity remained approximately constant throughout the cell cycle (Figure S7A), indicating that this marker is produced at a rate proportional to volume growth rate (Figure S7B), the expected behavior for many native proteins. In contrast, during osmotic oscillations, mCrimson fluorescence concentration increased, rising linearly to ~50% above normal levels after 4 h of 10-min, 500-mM oscillations (Figure 3A-C). When cells entered supergrowth, mCrimson intensity progressively decreased back to wild-type levels over 2-3 generations (Figure 3A-C), similar to the time scale for restoration of normal growth rates.

**Figure 3:**
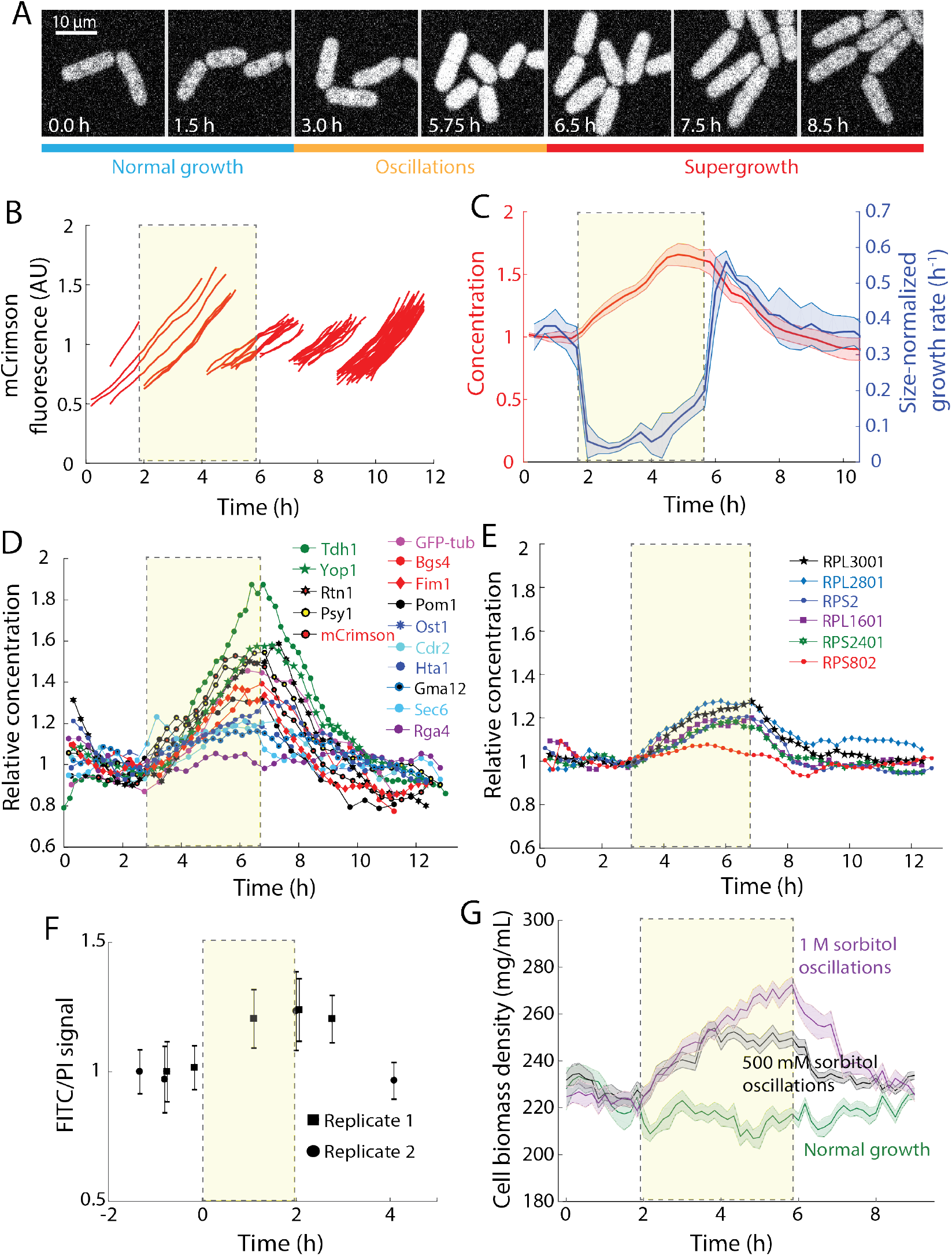
Osmotic-shock oscillations disrupt coupling between protein synthesis and cell size. A. Representative time-lapse images of mCrimson-expressing cells before, during, and after 24 10-min periods of 0.5-M sorbitol osmotic shocks. Fluorescence intensity increased during oscillations, and then gradually reverted to pre-shock levels during supergrowth.
B. Representative trajectories of integrated cellular fluorescence before, during, and after oscillations (n =54 cells). Synthesis of mCrimson continued relatively unabated during oscillations. The yellow box in this and subsequent panels denotes the 24 10-min periods of 0.5-M sorbitol osmotic shocks. (n = 54 cells)
C. mCrimson concentration increases by ~60% during oscillations (red), despite large decreases in volume growth rate (blue) (*n* = 843 cells). During supergrowth, the increased growth rate led to dilution of the mCrimson concentration. Dark centerline is the population-averaged mean, and the shading represents one standard deviation.
D. Concentrations of various proteins labeled with fluorescent proteins before, during, and after oscillations. All protein concentrations increased during oscillations, but to varying degrees.
E. Ribosomal proteins also increased during oscillations, but to a lesser degree; RPS802 concentration remained approximately constant throughout the experiment.
F. The ratio of FITC (proteins) to PI (DNA) fluorescence, a proxy for global protein density, increased during oscillations and then decreased back to steady-state levels during supergrowth. Cells were subjected to 12 cycles of 500-mM sorbitol shocks with 10-min period (Methods). Error bars are one standard deviation (n = 200 cells each datapoint).
G. Cytoplasmic density, as measured by quantitative phase imaging (Methods), increased during osmotic shocks and decreased during supergrowth. Dark centerline is the population-averaged mean cell density, and the shading represents one standard deviation. Green: normal growth (n = 88 cells); black, 500-mM oscillations (n = 187 cells); purple, 1-M oscillations (n = 106 cells).

To investigate why mCrimson accumulates during osmotic oscillations, we computed the rates of mCrimson production and compared to growth rate. During oscillations, the rate of mCrimson production remained close to normal levels even though cells grew in volume at a much slower rate than normal (Figure S7D). Thus, mCrimson increased in concentration not because of a large increase in protein production, but because of a failure to decrease protein production as much the reduction in volume growth rate.

We also examined how mCrimson concentration relaxed to equilibrium levels during supergrowth. In this phase, the cells grew in volume much faster than mCrimson production (Figure S7D). Much of this decrease in concentration can thus be explained by a dilution mechanism, which is enhanced by the rapid growth. The rate of “production” was also even lower than normal levels (Figure S7D), which may reflect decreased protein synthesis or increased degradation rates. Calculations using our growth-rate measurements to predict the decay constant of mCrimson fluorescence, assuming steady-state production after oscillations, indicated that at least 75% of the fall in mCrimson levels relative to controls can be explained by dilution during growth. These findings highlight dilution by rapid growth as a mechanism contributing to protein homeostasis.

Next, we determined whether native cellular proteins also increase in concentration during oscillations. We imaged fluorescently tagged proteins representing a variety of cellular processes, such as metabolism, chromosomal organization, cell-size control, and cell growth (Figure 3D, Table S1). These proteins all maintained a constant concentration during normal growth. Much like E2-mCrimson, the concentrations of many of these cellular proteins increased during 4 hours of 10-min, 0.5 M osmotic oscillations, and then returned gradually to normal during supergrowth (Figure 3D). No aggregates of these tagged proteins were observed. Similar changes in intensities were observed for fusions to various fluorescent proteins (Figure 3D), suggesting that the behavior was not an artifact of any particular fluorescent protein. Among the different protein fusions, there was a large range of behavioral variation, with different rates of increase and decrease during oscillations and supergrowth. Impressively, some proteins accumulated by ~100% during oscillations. In contrast, a subset of proteins displayed modest or no increases in concentration during oscillations. For example, the concentration of histone H2A was relatively steady throughout the experiment (Figure 3D), perhaps reflecting the fact that the amount of DNA is not increasing abnormally in these cells. The cell-size sensor Cdr2 also increased only modestly during oscillations (Figure 3D), consistent with the maintenance of size control. Ribosomal proteins also showed only modest increases (Figure 3E; see section below). These findings suggest that osmotic oscillations caused large changes in the concentrations of a significant subset of cellular proteins.

We next employed two complementary methods to measure the global state of cellular components. Staining of individual cells for total protein with fluorescein isothiocyanate (FITC) showed a similar rise and fall, with a mean 20% increase during 2 hours of osmotic oscillations (Figure 3F), while DNA concentration remained constant (Figure S8). Quantitative-phase imaging (Methods) showed that intracellular density, as measured from the diffractive index of the cell, also exhibited a ~10% and 20% increase during 0.5-M and 1-M oscillations, respectively (Figure 3G). Both measurements were in approximate agreement with the average increases in protein concentrations (Figure 3D). Thus, osmotic oscillations clearly produce global changes in protein concentration and cytoplasmic density.

Other oscillation regimes induced even larger increases in protein concentration. For instance, over the course of 48 oscillation cycles with 1 M sorbitol, cells halted growth and accumulated 150% higher mCrimson concentrations (Figure S9). These additional oscillatory cycles of 1M sorbitol displayed no sign of saturation in linear concentration accumulation (Figure S9B), suggesting that even higher increases are likely possible.

We sought to determine if a simple model of growth kinetics could quantitatively predict such a linear increase during growth inhibition. Experimental measurements have shown that translational efficiency 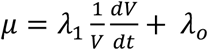 contains a growth-dependent and growth-independent component *λ*_0_^29^, which leads to a prediction of steady-state growth rate that is within 30% of experimental measurements (see Supplementary Information for more details). In our experiments with 1-M oscillatory shocks, growth halts completely and ribosome concentration is constant. In this scenario, our model predicts that the baseline translational efficiency *λ*_0_^29^ results in accumulation of biomass *M* at a constant rate *dM/dt* = *λ*_0_*R*, where *R* is the ribosome abundance. Hence, *M* is predicted to increase linearly during osmotic shock oscillations, at a reduced rate relative to unperturbed growth that yields a ~12% increase over 4 hours, similar to our experimental measurements (Figure 3G). Taken together, our experimental measurements and theoretical model indicate that oscillations and supergrowth represent periods of decoupling of protein production and cell expansion, and lead to profound accumulation of intracellular proteins.

### Increased protein concentrations drive supergrowth

Next, we asked whether the increased concentration of proteins is responsible for supergrowth. First, further analysis of our extensive exploration of oscillation parameters revealed that the maximal supergrowth rate was highly correlated with the observed increase in mCrimson concentration during oscillations (Figure 4A, *r* = 0.85). Second, we tested whether the increase of protein concentration is necessary for supergrowth. To globally inhibit protein synthesis, we treated cells with 100 μg/mL cycloheximide^30^ during oscillations, and then washed out the inhibitor upon exit from oscillations. These cycloheximide-treated cells recovered to near-normal growth rates but never achieved supergrowth (Figure 4B). Thus, efficient protein translation during oscillations is required for supergrowth.

**Figure 4:**
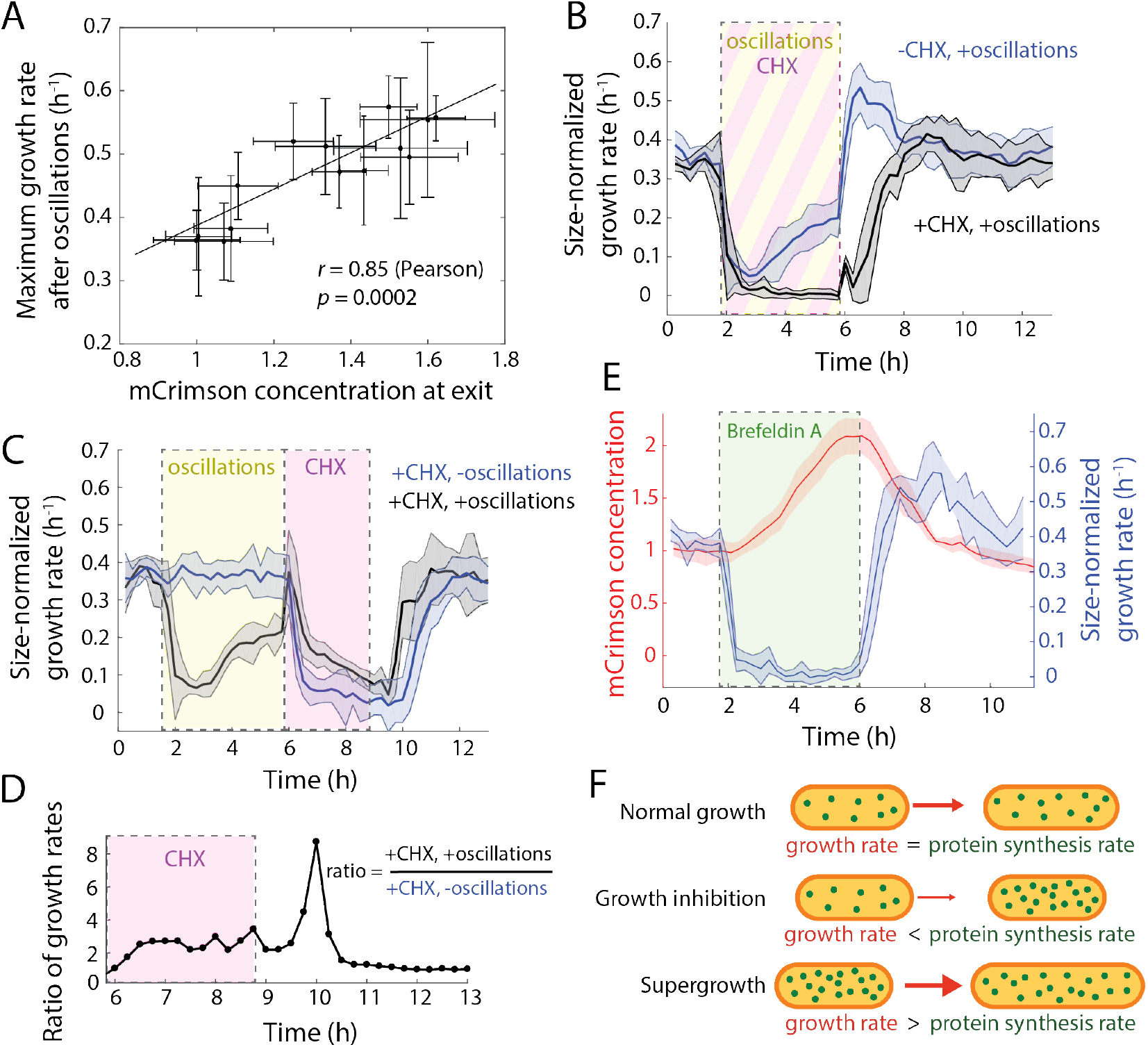
Excess protein concentration is necessary for supergrowth. A. Maximum size-normalized growth rates were highly correlated with the mean mCrimson concentration at exit from oscillations, across all shock magnitudes, periods, and number of shock cycles in Figure 2F. Error bars represent one standard deviation.
B. Cells treated with 100 μg/mL of the translation inhibitor cycloheximide (CHX; black shading, *n* = 228 cells) during oscillations (24 10-min periods, 0.5-M sorbitol shocks; yellow shading) rapidly halted growth during oscillations and exhibited no supergrowth afterward, unlike untreated cells (blue, *n* = 545 cells). Shading represents one standard deviation.
C. Cells exposed to osmotic-shock oscillations as in (B) (black, *n* = 175 cells) or maintained in constant YE5S growth conditions (blue, *n* = 260 cells) were treated (directly after the exit from oscillations) for 3 h with 100 μg/mL cycloheximide. Cells that underwent oscillations grew more during CHX treatment and resumed normal growth more quickly after CHX washout. Shading represents one standard deviation.
D. During cycloheximide treatment, the ratio of growth rates of cells that did and did not undergo oscillations was always ≥1 and reached a plateau within 30 min. At the plateau, cells that underwent oscillations grew approximately three-fold faster than control cells. The growth-rate ratio remained >1 for 2-3 h after cycloheximide treatment.
E. Brefeldin A (green shading) treatment completely inhibited growth, while mCrimson continued to accumulate throughout treatment. After the drug was washed out, cells initiated supergrowth with dynamics similar to those achieved after osmotic shocks.

Finally, we tested whether the accumulation of growth materials during oscillations is sufficient to drive growth in the absence of new protein synthesis. For control cells that were not exposed to oscillatory shocks, cycloheximide treatment caused a rapid cessation in cell growth. In contrast, cells that had been exposed to oscillations still grew substantially, at ~3-fold faster growth rates than control cells for 2 h (Figure 4C,D). Cells after oscillations also reestablished steady-state growth rates upon removal of cycloheximide more quickly (Figure 4C). Collectively, these results support the storage model in which cells with accumulated components are able to undergo some growth even in the absence of new protein synthesis.

### Effect of ribosomal concentration, cell wall synthases, and turgor pressure on supergrowth

To probe whether the up-regulation of particular cellular processes drives supergrowth, we assessed three candidates involved in growth regulation: ribosomes, the cell-wall biosynthetic machinery, and turgor pressure. We monitored ribosomal protein abundances in single cells using ribosomal protein-GFP fusions expressed from their endogenous promoters (Methods) and found that their steady-state concentrations were relatively constant (Figure 3E). The concentrations of these ribosomal proteins increased only modestly during oscillations (from 0−20%) (Figure 3E) and then quickly reverted to baseline levels during supergrowth. Some ribosomal proteins displayed nearly constant concentration throughout the experiment (Figure 3E, S10), indicating that their production rates decreased and increased concurrently with the growth rate. Thus, rapid supergrowth is not caused by a large increase in ribosome concentration.

As cell-wall assembly likely limits the rate of cell expansion, a candidate factor for controlling cell-wall growth is the major cell-wall glucan synthase Bgs4. Bgs4 is a trans-membrane protein that localizes to growing cell tips during interphase and to the developing septum during cytokinesis^31^. During the oscillations, as cells grew slowly (Figure S11A), Bgs4 puncta were delocalized across the cell membrane (Figure S11B), consistent with the general depolarization of actin and Cdc42 seen in the osmotic-shock response^31–33^. Bgs4 intensities at cell tips decreased by 50%, although total Bgs4 concentration gradually increased (by ~50%) during this period (Figure 3D). Upon exit from oscillations, Bgs4 rapidly repolarized to the cell tip, and tip intensities rose to more than 50% more than in control cells within 30-45 min (Figure S11B). The amount of Bgs4 at the cell tips strongly correlated with the growth rate in these experiments (*r* = 0.95) (Figure S11C). It is likely that such increases in the levels of Bgs4 and additional polarity factors contributing to cell wall growth collectively contribute to rapid growth. These results reveal mechanistic insight into why cell growth slows down during osmotic shock oscillations and how polarized growth is rapidly established after oscillations.

Another potential contributor to supergrowth could be an increase in turgor pressure that may arise from increased cytoplasmic density. We examined turgor pressure by measuring cell width, as changes in turgor pressure are expected to lead to concurrent reversible swelling in cell width independent of tip elongation. Although cell widths increased slightly during osmotic oscillations, they did not decrease during supergrowth as cytoplasmic density decreases (Figure S5A, S12). These data indicate that the small width increases during oscillations are not reversible and thus may arise from delocalized cell wall synthesis (Figure S11B,C)^32^, rather than from increases in turgor pressure. In sum, turgor pressure is unlikely to increase substantially during supergrowth, and is not the driver of the increase in growth rate during supergrowth.

### Transient inhibition of protein secretion also drives subsequent supergrowth

Our model predicted that other perturbations that transiently inhibit cell volume growth would also lead to proteome accumulation and subsequent supergrowth. One way of stopping growth and increasing buoyant density is through inhibition of secretion, as shown initially in budding yeast^34^. To halt secretion, we treated mCrimson-expressing cells for 4 hours with 100 μg/mL brefeldin A (BFA), which inhibits protein transport from the endoplasmic reticulum to the Golgi apparatus^35,36^. Cells exhibited a rapid decrease in growth rate to essentially zero throughout BFA treatment, during which time mCrimson concentration increased approximately linearly (Figure 4E). Many cells eventually died, but the 45% of cells that survived the treatment exhibited supergrowth upon BFA wash out (Figure 4E) with growth-rate trajectories similar to those observed after 0.5 M sorbitol oscillatory osmotic shocks (Figure 2D). Thus, rather than just a response to osmotic stress, supergrowth and proteome accumulation can be produced by other treatments that limit volume expansion without proportional inhibition of protein synthesis.

## DISCUSSION

Here, our interrogation of the growth response of *S. pombe* cells to osmotic perturbations has revealed unexpected insights into the relationships between cell size, growth, and biosynthesis. In investigating the basis for supergrowth, we find that these unprecedented rapid growth rates arise from a decoupling of biosynthesis and volume expansion. During the osmotic oscillations, volume expansion is impaired far more than biosynthesis, which leads to global accumulation of cellular components inside the cell. Release from these conditions then results in abnormally rapid growth that drives an accelerated return to normal proteome concentrations relative to dilution. Our results thus support a model in which rapid growth is driven by the accumulation of cellular materials. This effect of material storage on growth rate may be connected with the mechanisms underlying the near-universal scaling of growth rate with cell size (exponential growth) observed under steady-state environmental conditions across the kingdoms of life. In the simplest interpretation, cells in supergrowth grow rapidly because they have the internal resources of a larger cell. Whether the cellular materials that promote growth are some subset of the cytoplasmic constituents, or the entire collection, remains to be seen. Our data suggests that growth rate is set by a combination of cellular processes, including the biosynthetic machinery, metabolism, and cell-wall assembly, rather than by a single component such as ribosomes.

This work reveals that relatively simple manipulations can decouple cell growth and protein synthesis, yielding mechanistic insights into how these fundamental processes are normally coordinated. As concentrations of many proteins do not vary greatly during steady-state growth, it has been speculated that extensive feedback mechanisms exist for protein homeostasis; for instance, increased concentration of a certain protein may inhibit its own translation or transcription through a signaling pathway that senses its concentration, leading to a decrease back to normal levels; such a protein-specific mechanism has been suggested for tubulin homeostasis for instance^37^. In this scenario, we speculate that osmotic oscillations lead to breakdown of the putative feedback mechanisms, possibly because of the rapid changes in cell size and concentrations. However, our findings also raise a second, simpler possibility in which signaling feedback mechanisms may not exist for many proteins^38–40^. We speculate that volume expansion itself could act as a homeostatic mechanism, wherein biomass increases largely independent of volume and small variations in cytoplasmic protein concentrations are brought back into balance by a correction in short-term volume growth rate. Abnormal increases in protein concentrations would drive faster growth than synthesis, which would decrease their concentration in part through dilution, similar to what we observe during supergrowth. A decrease in protein concentration would lead to slowed growth, allowing protein concentration to build back up. Our experiments were able to reveal such a mechanism for the coordination of growth and volume, because they allowed us to uncouple this coordination and examine the recovery process.

Such a mechanism however may apply to a subset of proteins, but not to all. During osmotic oscillations, certain groups of proteins (such as metabolic enzymes) maintained relatively high synthesis rates and hence rose in concentration. By contrast, other proteins – including ribosomal proteins, histones, and cell size regulators – are likely regulated so that their expression scales with growth to maintain their concentrations; the level of accumulation of these proteins during osmotic oscillations would then reflect the robustness of this regulation. Tight control of ribosomal protein concentration is predicted to be another factor to keep the proteome in check^41^, preventing accumulation at more rapid rates when expansion is perturbed.

The remarkable cellular phenotype we have presented in which the concentrations of much of the proteome are altered, accompanied by substantial increases in cytoplasmic density that likely involve a sizeable replacement of water within the cytoplasm by protein biomass. Nevertheless, *S. pombe* cells retain viability, grow with normal morphologies, and exhibit normal periods of mitosis and cytokinesis, underscoring the robust control of critical biochemical processes. In particular, this work provides a striking demonstration of how cell size can be maintained even over a 10-fold range of growth rates, a finding inconsistent with models of cell-size control based directly on growth rate^42^. Rather, these results support the existence of cell-size control mechanisms in fission yeast cells that involve direct assessment of cell size through “sizers”^43,44^. In general, our findings indicate that cellular density can be regulated or perturbed in various settings such as stress. Procedures such as osmotic oscillations that increase protein expression per unit volume may have practical applications, for instance in protein production. It will be valuable to explore in future studies the myriad ways in which such global changes in cytoplasmic density affect cellular processes.

## METHODS

### Strains and growth conditions

All strains used in this study are listed in Table S1.

Single colonies were inoculated from plates into liquid rich medium (YE5S, Sunrise Science Products YES-225) overnight with shaking at 30°C. Once cells reached mid-log phase (OD_600_~0.5), they were diluted 1:10 directly into ONIX Y04C-02 microfluidic flow cells (EMD Millipore) that had been primed with fresh medium for 15 min. Cells were grown for 15-30 min prior to imaging to allow them to equilibrate to the flow-cell chamber. YE5S was supplemented with D-sorbitol (Sigma) for hypertonic conditions, and fresh medium was introduced into the culture chambers at 5 psi. Control measurements were performed by exchanging YE5S between wells in adjacent culture chambers. By adding a tracer dye (0.5 μg/mL Alexa Fluor 647 carboxylic acid; Life Technologies) to the sorbitol-supplemented medium, we determined that media exchange occurred within 10 s (Figure 2A, S3A). Where applicable, drug (cycloheximide or brefeldin A; Sigma) was added to the medium.

### Microscopy

Cells were imaged in phase-contrast using a Ti-Eclipse stand (Nikon Instruments) with a 40X (NA: 0.95) or 100X (NA: 1.45) objective, and a Zyla 4.2 sCMOS camera (Andor Technology). Temperature was maintained with a stage-top incubator (OkoLab), which was warmed for at least 1 h prior to imaging. Images were acquired at 1-s or 5-s frame intervals for measuring rapid volume changes during osmotic shocks (Figure S3B), and 1-min or 5-min frame intervals for osmotic shock and growth measurements. For strains expressing fluorescent proteins, cells were imaged in phase-contrast and with laser illumination in spinning-disk confocal mode (Yokogawa CSU-10), and images were acquired with an EM-CCD camera (Hamamatsu) every 10 min at low power to avoid bleaching. The microscope systems were integrated using μManager v. 1.41^45^.

### Cell segmentation and size/fluorescence quantification

We first utilized the deep neural network-based machine learning framework *DeepCell* (Van Valen et al., 2017) to segment cells. For each imaging technique (wide-field, confocal) and objective, approximately 200 cells were manually outlined to produce a training dataset. Trained networks were used to generate binary images for feature (extracellular/cell perimeter/cytoplasm) identification. These images were used as the input for gradient segmentation in *Morphometrics* v. 1.1^46^ to define cell contours at sub-pixel resolution.

The centerline between the two poles was calculated through an iterative method in which symmetric bisections were created starting from the contour’s centroid toward the poles. Cell length was defined as the total length of the centerline, and width as the median length of lines running perpendicular to the centerline and stretching between the two sides of the cell. Cell surface area and volume were calculated by integrating disks of revolution at each point on the centerline.

For fluorescence quantification, the total signal within each cell contour was summed.

### Estimation of supergrowth contribution to mCrimson dilution

Estimates for growth-based dilution during supergrowth were calculated using the equation

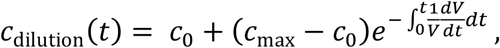

where the rate of protein production during supergrowth is assumed to follow the growth rate and 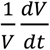 is the population-averaged growth rate. The difference between the actual and estimated concentrations (*c*(*t*) − *c*_dilution_(*t*)) was used to estimate the contribution to dilution from growth at >75%.

### Bgs4 tip localization and intensity quantification

Cell tips were identified based on peaks in the contour curvature, which is defined at each contour point as the second derivative of the local vector normal. The fluorescence intensity was integrated along a line scan perpendicular to the contour at each contour point whose curvature was greater than 0.12 μm^−1^, a threshold that was determined visually as an optimal identifier of cell tip regions.

### Osmotic oscillations on bulk cultures

Using a tabletop upright glass vacuum filter holder (Fisherbrand) with 0.45-μm HA membrane filters (Millipore, HAWP04700), the medium was rapidly exchanged (<30 s) for cultures of up to 200 mL. All media were maintained at 30°C, and cells were resuspended and shaken at 30°C during each half-period. One milliliter samples were taken approximately each hour during the experiment and immediately fixed in 70% cold ethanol. Cells from the culture were imaged directly after these oscillations to measure growth rate and mCrimson fluorescence.

### Fluorescein isothiocyanate (FITC) and propidium iodide (PI) staining

Staining protocols were adapted from Ref. ^47^. One-milliliter samples were collected at various time points during the bulk oscillations protocol at OD600 ~ 0.5, and cells were fixed with 70% cold ethanol for at least 24 h at 4°C. Approximately 10^6^ cells (300 μL) were washed with 50 mM sodium citrate buffer (pH 7.4), and then treated with 0.1 mg/mL RNAse A (ThermoFisher, EN0531) for at least 2 h. For total protein staining, cells were washed and then stained with 5 ng/mL FITC (Sigma, F7250) for at least 30 min, then washed three times. To determine total DNA content, cells were stained with PI at 0.1 mg/mL for at least 30 min and then washed.

To quantify FITC/PI ratios, cells were imaged on 1% agarose + EMM pads (Figure S8). *Z*-stacks were acquired to obtain sum-projections after background subtraction, with cell outlines obtained manually due to the poor phase contrast of fixed cells.

### Quantitative Phase Imaging

Images were acquired with a Ti-Eclipse Ti inverted microscope (Nikon) equipped with a 680-nm bandpass filter (D680/30, Chroma Technology) in the illumination path with a 60X (NA: 1.4) DIC oil objective (Nikon). Before imaging, Koehler illumination was configured and the peak illumination intensity with 10-ms exposure time was set to the middle of the dynamic range of the Zyla sCMOS 4.2 camera (Andor). μManager v. 1.41^45^ was used to automate acquisition of brightfield z-stacks with a step size of 250 nm from ±3 μm around the focal plane (total of 25 imaging planes) at 10-min intervals.

For analysis, all imaging planes between maximal distances of +1.25 μm and −1.25 μm around the focal plane were selected. Based on these images, the phase information was calculated using a custom Matlab script implementing a previously published algorithm^48^. In brief, this method relates the phase information of the cell to intensity changes along the z-direction. Equidistant, out-of-focus images above and below the focal plane are used to estimate intensity changes at various defocus distances. A phase-shift map is reconstructed in a non-linear, iterative fashion to solve the transport-of-intensity equation.

A Gaussian peak was fitted to the background of each image and then corrected to be at zero phase shift using an image-wide subtraction of the mean of the peak. These corrected images were segmented using *DeepCell* and *Morphometrics* v. 1.1. From the cell outlines, the median intensity of each cell was obtained with ImageJ and used to calculate the cytoplasmic density as follows. The difference of the reconstructed phase map median intensity of cells in YE5S medium compared to cells in YE5S medium supplemented with 100 mg/mL BSA was used to define the phase shift contribution equivalent to 100 mg/mL biomass. The phase shift corresponding to 100 mg/mL BSA was then calculated by first averaging the intensity of all cells in YE5S medium at timepoints 10 min before and after BSA addition. Then, the difference relative to the median intensity of all cells during the measurement in YE5S supplemented with 100 mg/mL BSA was defined as the signal contributed by 100 mg/mL biomass.

Throughout the experiment, multiple measurements in YE5S supplemented with 100 mg/mL BSA were performed and calibration values for each imaging timepoint in between two consecutive measurements in YE5S + BSA were obtained by linear interpolation. These values were then used for quantification of cell density.

## ACKNOWLEDGMENTS

We thank members of the Huang, Chang, J. Skotheim, and S. Dumont labs for helpful discussions. This work was funded by NIH CAREER Award MCB-1149328 (to K.C.H.), the Allen Discovery Center at Stanford University on Systems Modeling of Infection (to K.C.H.), the National Natural Science Foundation of China (NSFC; 3167080548) (to X.H.), and NIH R01-GM56836 and Bilateral BBSRC-NSF/BIO Award 1638195 (to F.C.). K.C.H. is a Chan Zuckerberg Biohub Investigator.

## AUTHOR CONTRIBUTIONS

B.D.K., K.C.H., and F.C. conceptualized the study. B.D.K., P.O., E.R.R., X.H., K.C.H., and F.C. designed the experiments. B.D.K. and P.O. performed oscillatory osmotic shock experiments. P.O. performed quantitative phase imaging. X.X. and W.C. constructed ribosomal-GFP strains. B.D.K., P.O., E.R.R., K.C.H., and F.C. analyzed data. B.K., K.C.H., and F.C. wrote the manuscript. All authors reviewed the manuscript before submission.

## AUTHOR INFORMATION

Reprints and permissions information is available at www.nature.com/reprints. The authors declare no competing interests.

