## Supplement for "Decoupling of rates of protein synthesis from cell expansion leads to supergrowth"

**Supplementary Information for "Decoupling of rates of protein synthesis from cell expansion leads to supergrowth"**

443 Via Ortega

Stanford, CA 94305-4125 USA

### Supplementary Text

Below, we describe an analytical model for the increase in proteome concentration during osmotic shock oscillations.

#### ***Setup of model based on strict regulation of ribosome concentration***

In our oscillatory osmotic-shock experiments, ribosomal protein concentration was relatively constant compared to non-ribosomal proteins. Imposing the requirement of constant ribosome ribosome concentration $\rho_R = R/V$, where $R$ is the number of ribosomes and $V$ is the cell volume,

$$\frac{d\rho_R}{dt} = 0. \tag{1}$$

Eq. 1 imposes that $R$ and $V$ are related by

$$\frac{1}{R}\frac{dR}{dt} = \frac{1}{V}\frac{dV}{dt}. \tag{2}$$

During growth, biomass $M$ is converted into irreversible volume expansion at a rate $\alpha$[1], such that

$$\frac{dV}{dt} = \alpha M, \tag{3}$$

while we assume that the biomass production rate is approximately set by the global protein synthesis rate $\mu$:

$$\frac{dM}{dt} = \mu R. \tag{4}$$

We also know from previous studies that the efficiency of translation has a linear growth rate dependence

$$\mu = \lambda_1 \frac{1}{V}\frac{dV}{dt} + \lambda_0, \tag{5}$$

And we fitted the data in Ref. [2] to obtain values of $\lambda_1 \approx 14 \frac{\frac{\text{aa}}{\text{s}}}{\text{ribosome}} \frac{1}{\text{h}^{-1}}$ and $\lambda_0 \approx 0.72 \frac{\frac{\text{aa}}{\text{s}}}{\text{ribosome}}$ as measured in amino acids produced (aa).

***Predictions for steady-state dynamics***

To solve the coupled set of first-order differential equations in Equations 4 and 5, we can write

$$\frac{dM}{dt} = \left(\lambda_1 \frac{1}{V}\frac{dV}{dt} + \lambda_o\right) R = \lambda_1 \rho_R \frac{dV}{dt} + \lambda_o R.$$

Finally, using Equation 3,

$$\frac{dM}{dt} = \lambda_1 \rho_R \alpha M + \lambda_o R. \tag{6}$$

During normal (rapid) growth, we can ignore the second term during normal growth since it is roughly 10-fold smaller than the first term, giving

$$\frac{1}{M}\frac{dM}{dt} \approx \lambda_1 \rho_R \alpha.$$

At steady state, $\frac{1}{M}\frac{dM}{dt} = \frac{1}{V}\frac{dV}{dt}$, thus the growth rate is a constant

$$\frac{1}{V}\frac{dV}{dt} = \lambda_1 \rho_R \alpha. \tag{7}$$

Using measured values, this equation predicts a growth rate of ~ 0.25 $h^{-1}$ [3], in reasonable agreement with our measurements of ~0.35 $h^{-1}$ in rich media at 30 °C. Furthermore, Equation 7 agrees with measurements showing that steady-state growth rate depends linearly on the ribosome concentration[1].

***Modeling of growth arrest dynamics***

During growth arrest, $\alpha = 0$, so volume growth rate becomes zero and the ribosome number remains constant: $\frac{dV}{dt} = 0$ and $\frac{dR}{dt} = 0$. Then, the biomass production rate is

$$\left.\frac{dM}{dt}\right|_{\alpha=0} = \lambda_0 R_0, \tag{8}$$

where $\lambda_0$ is the translational efficiency of the cell under growth arrest. Equation 8 agrees with the observation that protein concentration, as measured by fluorescence (Fig. 3C,F) and biomass density (Fig. 3G), increases linearly during osmotic shock oscillations, at a reduced rate relative to normal growth.

By dividing Equation 8 by the static volume under growth arrest, the biomass density, $\rho_B$, evolves as

$$\frac{d\rho_B}{dt} = \lambda_0 \rho_R.$$

Since $\lambda_0 \rho_R$ is a constant, we can integrate over a time interval, $\Delta t$, to obtain

$$\Delta\rho_B = \lambda_0 \rho_R \Delta t.$$

From Ref. [3], $\rho_R \approx 14000$ ribosomes/μm$^3$, and using the fitted value of $\lambda_0$ in Equation 5, a time interval of $\Delta t = 4$ h yields

$$\Delta\rho_B \approx 0.026\,\frac{\mathrm{pg}}{\mu\mathrm{m}^3}.$$

where an amino acid is $\sim 1.82 \times 10^{-10}$ pg. $\Delta\rho_B$ is the estimated contribution of protein mass to the cell's total biomass after 4 h of volume growth arrest.

The average yeast cell is ~1.1 g/mL[4], with ~20% of the cell's volume estimated to be occupied by biomass[5,6]. Using this estimate of the biomass density ($\rho_B^0 \approx 0.22\,\frac{\mathrm{pg}}{\mu\mathrm{m}^3}$), we obtain an increase of $\Delta\rho_B \sim 12\%$, in reasonable agreement with the biomass density increase of ~20% that we observed during 4 h of 1-M sorbitol oscillations (Figure 3G).

### Supplemental Figures

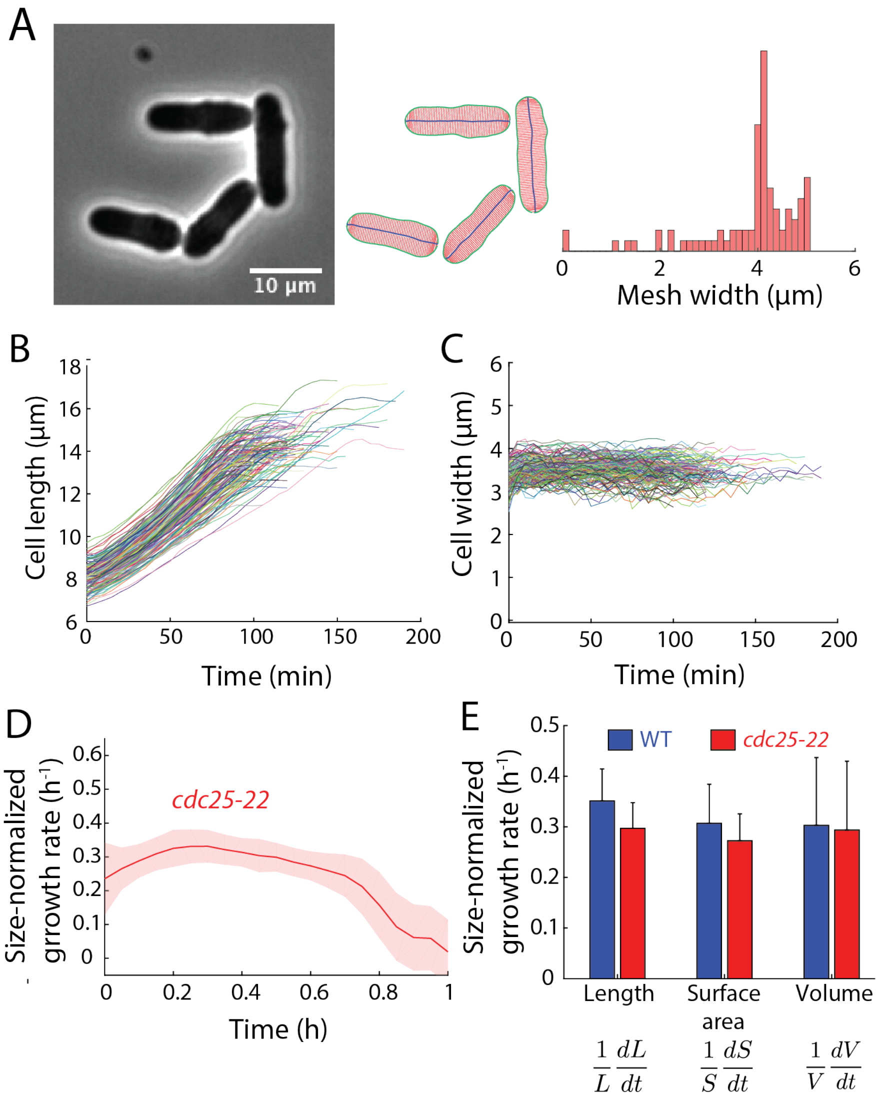


**Figure S1: *S. pombe* cells exhibit exponential growth for a large fraction of the cell cycle.**

A) Example phase-contrast image of cells grown in microfluidic chambers (left) and skeleton (middle; red: width, green: outline, blue: centerline) calculated from automated segmentation analysis (Methods). Right: the distribution of mesh widths of an example skeleton, with the overall cell width defined as the median of this distribution.

B) Cell length traces of a population of wild-type cells during steady-state growth in a microfluidic flow cell ($n$ = 147 cells).

C) Cell width traces of the population of wild-type cells shown in (B).

D) Elongation rate ($1/L\ dL/dt$) of *cdc25-22* cells during steady-state growth, excluding mitosis ($n$ = 246 cells).

E) Slopes of instantaneous rates ($dX/dt$ vs. $X$) as measured from length ($L$), surface area ($S$), or volume ($V$).

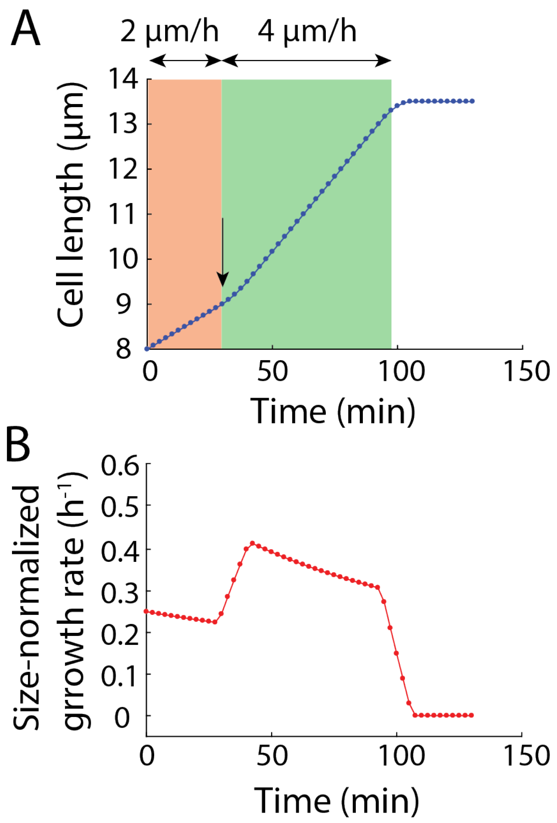


**Figure S2: Bi-linear growth model is inconsistent with experimental data.**

A) Simulation of bilinear growth according to growth rates and transition times proposed in the literature[7-10]. Cells were assumed to elongate at 2 μm/h for the first third of G2 and then elongation rate doubled to 4 μm/h until starting mitosis at 14 μm.

B) The size-normalized growth rate of a cell undergoing bilinear growth is clearly incongruent with our experimental measurements (Fig. 1D).

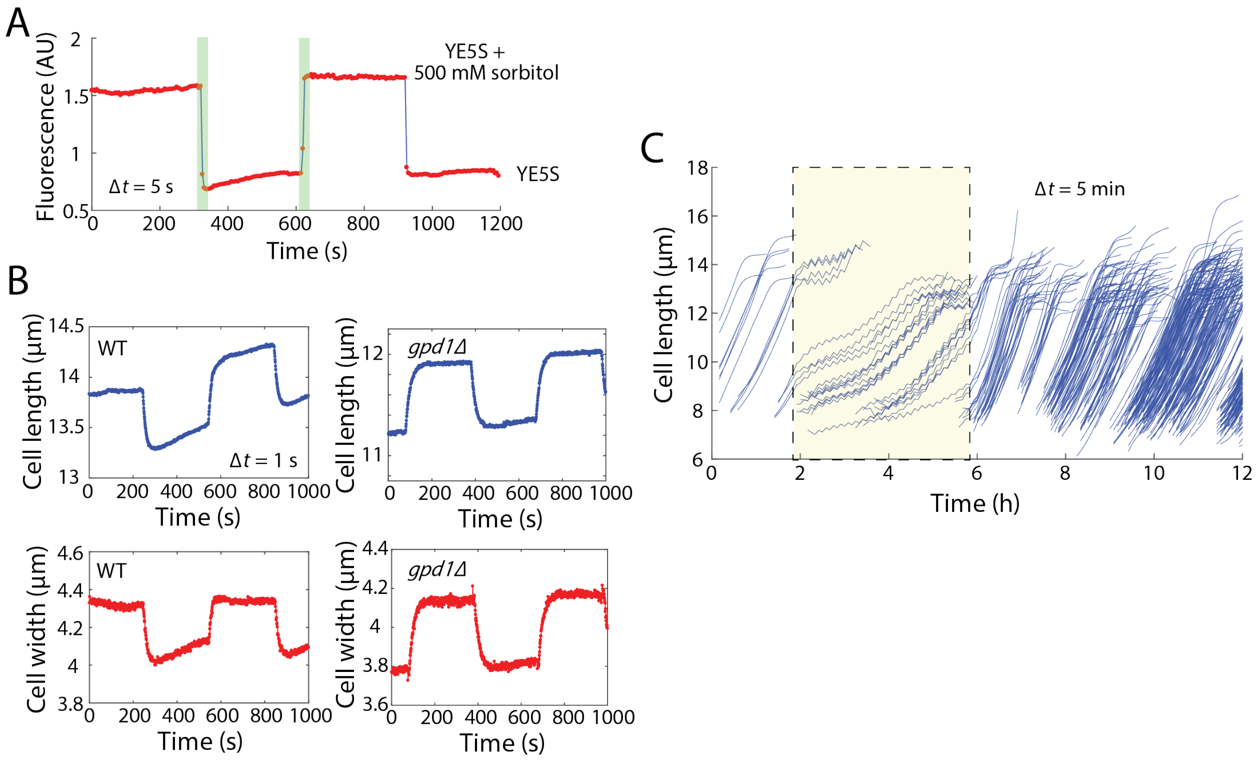


**Figure S3: Cell dimensions during oscillatory osmotic shocks**

A) Rich medium supplemented with 0.5 M D-sorbitol was tracked during oscillations (10-min period) by including a tracer dye (0.5 μg/mL Alexa Fluor 647 carboxylic acid). Fluorescence images (YFP) acquired every 5 s demonstrate that the media were exchanged in less than 10 s.

B) Length (blue) and width (red) of representative wild-type (left) and *gpd1Δ* (right) cells during oscillatory 0.5 M sorbitol osmotic shocks with 10-min period. Images were acquired every 1 s.

C) Length trajectories of wild-type cells undergoing oscillatory osmotic shocks. Yellow outlined box represents the 4-h duration of 0.5 M D-sorbitol shocks with 10-min period. Images were acquired every 5 min ($n$ = 302 cells).

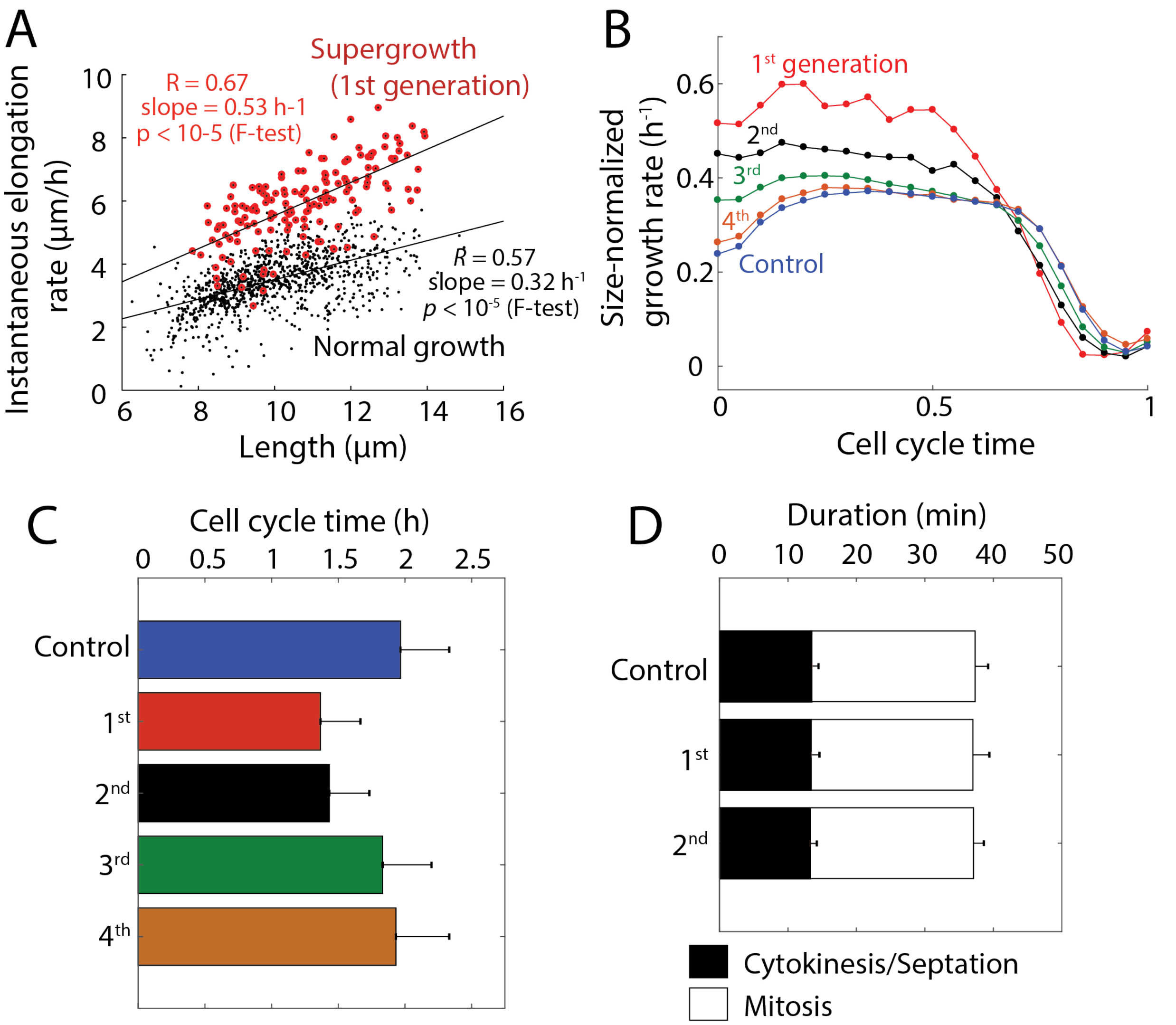


**Figure S4: Growth rate increases during supergrowth diminish each generation, and shorten G2 phase only.**

A) Instantaneous elongation rate increases with cell length during the first generation of supergrowth (red circles; *n* = 159). The slope of the best linear fit is steeper than that of normal growth (black dots; *n* = 998), indicating a higher size-normalized growth rate.

B) Size-normalized growth rates as a function of normalized cell-cycle time during each generation of supergrowth after 4-h of oscillatory 0.5 M D-sorbitol osmotic shocks with 10-min period. The first generation of supergrowth represents cells that divided within 15 min of exit from oscillations (*n* = 10 cells). Subsequent generations were classified based on manual inspection of trajectories. The second generation of supergrowing cells did not display the initial phase of super-exponential growth (Fig. 1d). Number of cells *n* (generation): 146 (2nd), 194 (3rd), 298 (4th), 929 (control).

C) Mean cell-cycle duration (including mitosis) increased during each generation of supergrowth after osmotic shock oscillations. Number of cells as in (B).

D) Duration of mitosis and cytokinesis were constant during supergrowth. Images were acquired every 1 min, and mitosis and cytokinesis were defined based on the onset of growth arrest and completion of septation, respectively. Number of cells (generation): 31 (1st), 34 (2nd), 53 (control).

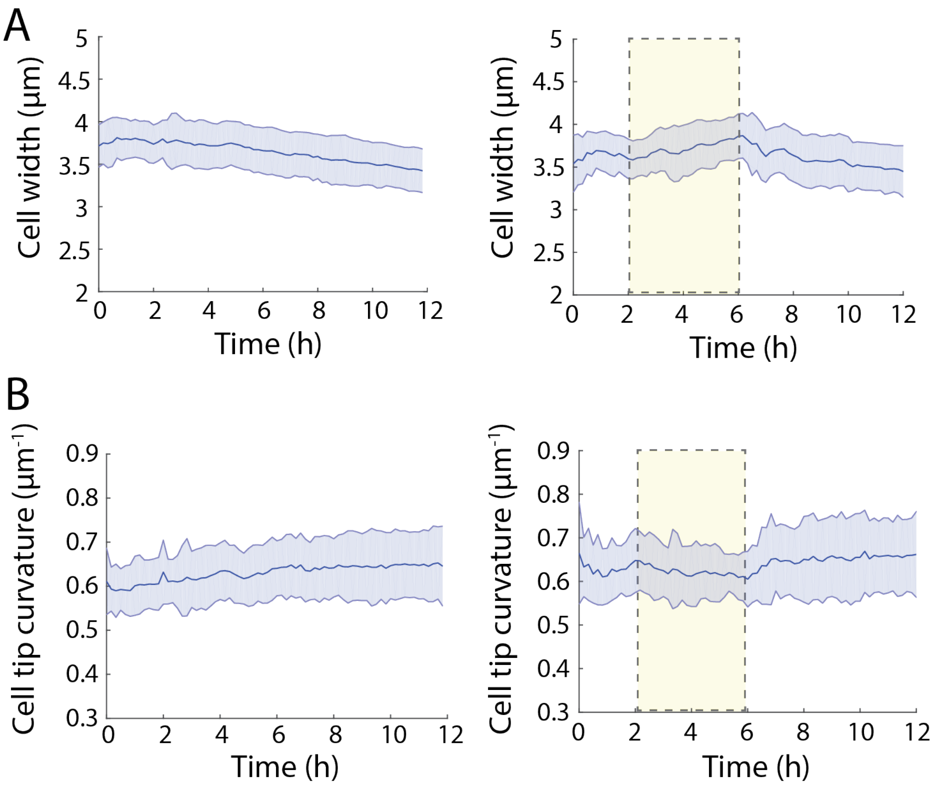


**Figure S5: Cell width and tip curvature remain mostly unchanged by osmotic shock oscillations and supergrowth.**

A) Cell width measurements during normal growth (left) and osmotic shock oscillations (right; yellow box represents 4 h of 0.5-M sorbitol oscillations with 10-min period). Apparent decreases in width during normal growth are due to cell crowding effects on automatic segmentation. Dark centerline is the population-averaged mean width and the shading represents 1 standard deviation (S.D.) (control, *n* = 1242 cells, control; oscillations, *n* = 973 cells).

B) Cell tip curvature measurements during normal growth (left) and osmotic shock oscillations (right; yellow box represents 4 h of 0.5-M sorbitol oscillations with 10-min period over 4 h). Dark centerline is the population-averaged mean cell tip curvature and the shading represents 1 S.D. (control, $n$ = 1242 cells; oscillations, $n$ = 973 cells).

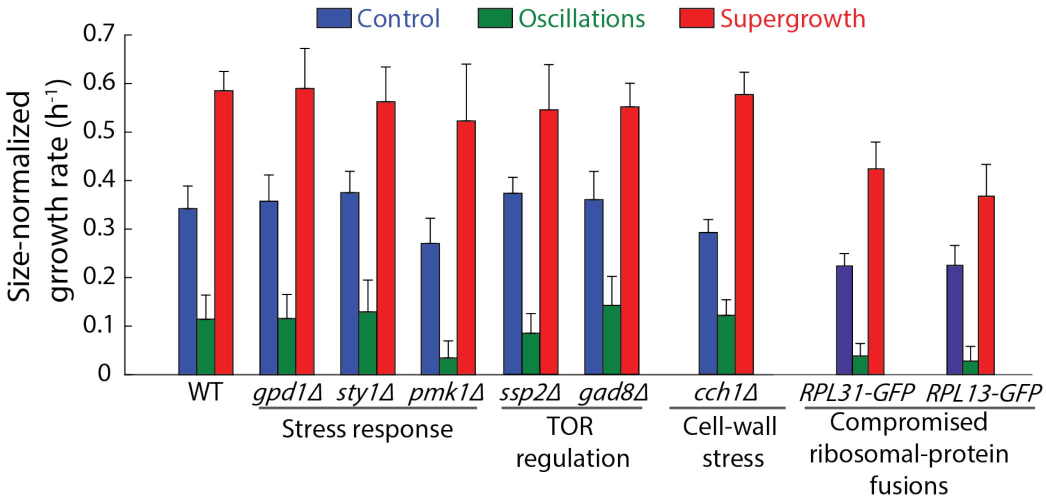


**Figure S6: Osmotic stress response and growth mutants exhibit supergrowth behavior.**

Mutants related to stress response pathways and growth have similar responses to oscillatory osmotic shocks as wild-type (WT) cells. Control and oscillatory elongation rates were computed as the mean size-normalized growth rates during steady-state growth prior to osmotic shocks and during the 4 h of oscillations, respectively; supergrowth rate was defined as the peak in the growth rate after exit from oscillations, smoothed over a 15-min time window. Error bars represent 1 S.D. Number of cells: WT, *n* = 929; *gpd1Δ*, *n* = 641; *sty1Δ*, *n* = 395; *pmk1Δ*, *n* = 421; *ssp2Δ*, *n* = 443; *gad8Δ*, *n* = 634; *cch1Δ*, *n* = 197; *RPL31-GFP*, *n* = 99; *RPL13-GFP*, *n* = 62.

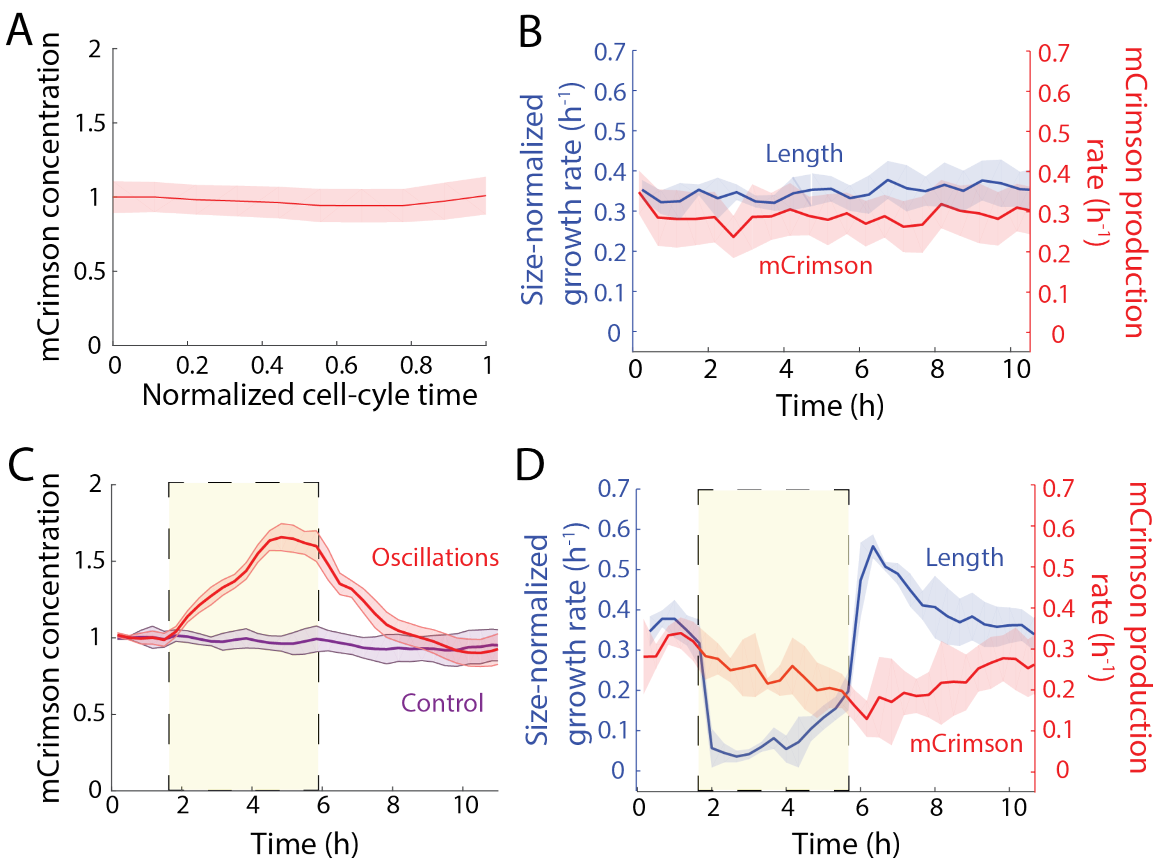


**Figure S7: Decoupling of mCrimson production from elongation leads to increase in mCrimson concentration.**

A) mCrimson concentration was approximately constant throughout the cell cycle during steady-state growth ($n$ = 254 cells). $t$ = 0 represents the birth time, and time is normalized to the time between birth and division (as defined by 'snapping'). Dark line is the population mean and shaded bars represent 1 S.D.

B) Population-averaged mCrimson production rate and size-normalized growth rate were approximately constant during steady-state growth (excluding mitosis). Fluorescence production rate was defined as the instantaneous cellular change in the log of the integrated intracellular fluorescence ($n$ = 254 cells).

C) Population-averaged intracellular concentration of mCrimson during growth (excluding mitosis). Cells undergoing osmotic shock oscillations (red) increased their mCrimson concentrations by ~60% relative to control cells (purple) over the course of the oscillations. Yellow outlined box represents the 4-h duration of 0.5-M sorbitol osmotic shocks with 10-min period. Control cells were grown in a chamber of the same microfluidic flow cell adjacent to cells exposed to oscillations ($n$ = 842 cells).

D) Population-averaged mCrimson production rate and elongation rate (excluding mitosis) during an oscillatory osmotic shock experiment. Yellow outlined box represents the 4-h duration of 0.5 M D-sorbitol shocks with 10-min period ($n$ = 842 cells).

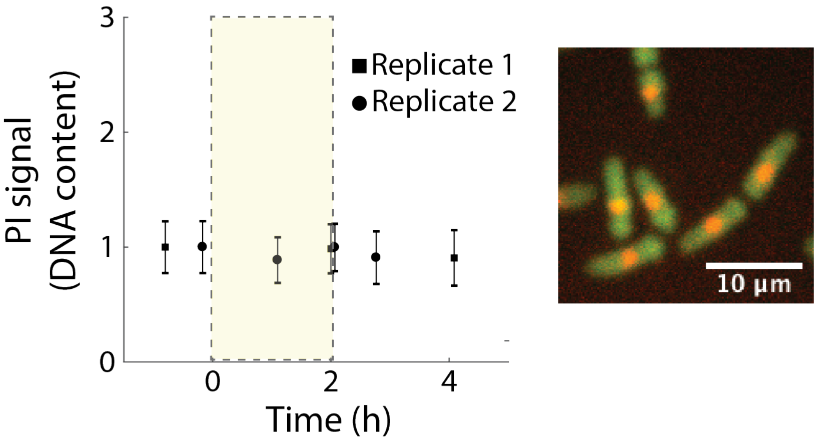


**Figure S8: DNA content does not change during osmotic shock oscillations or supergrowth**

Propidium iodide (PI) staining, which was used to estimate cellular DNA content (Methods), remained constant throughout both osmotic shock oscillations and supergrowth in non-mitotic cells. The yellow outlined box indicates 12 cycles of 500 mM sorbitol shocks with 10-min period (Methods; bulk oscillations). Error bars are 1 S.D. (*n* > 2000 cells).

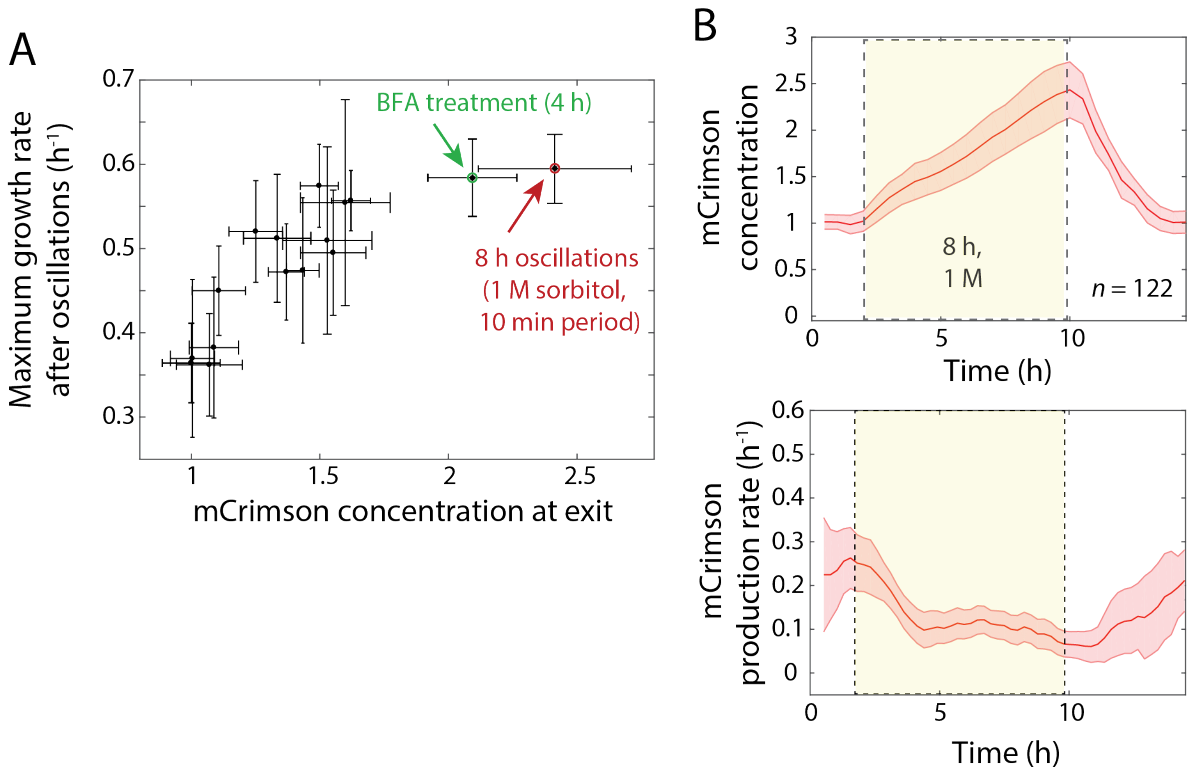


**Figure S9: Maximum supergrowth rate eventually saturates for long durations of osmotic shock oscillations.**

A) Maximal supergrowth rate after osmotic shock oscillations as a function of mCrimson concentration at exit from oscillations. Data includes those from Fig. 4A, along with one point representing long-term (8 h), large-amplitude (1 M) osmotic shocks (red circle) and one point representing BFA-treated cells (green circle, Fig. 4E). The highest supergrowth rate achieved was ~0.6 $h^{-1}$ ($n$ = 2714 cells).

B) Top: concentration of mCrimson during an osmotic shock oscillation experiment over a long duration (8 h) of large-amplitude (1 M) shocks with 10-min period. Bottom: After an initial decrease over the first 2 h, mCrimson production stabilized at a rate ~0.1 $h^{-1}$ and persisted throughout the rest of the oscillations. The centerline is the population average, and shaded error bars are 1 S.D. ($n$ = 122 cells).

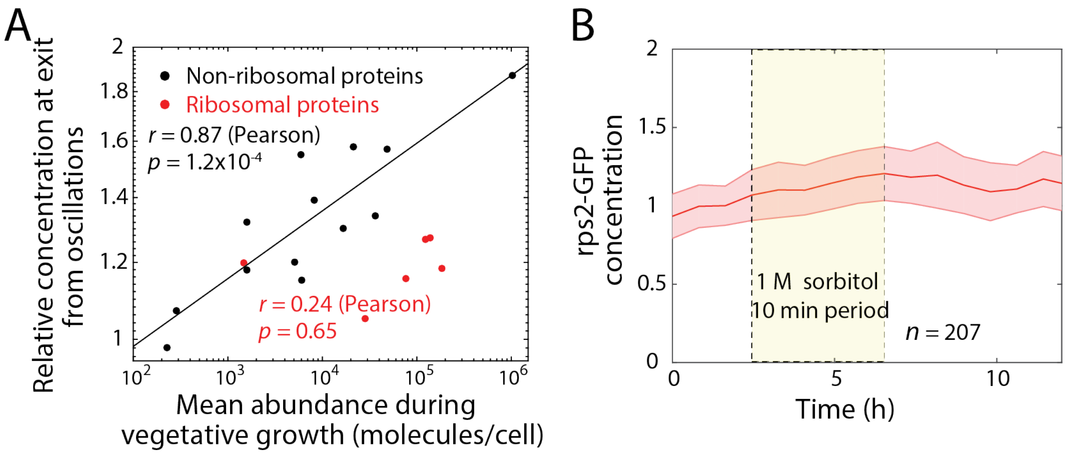


**Figure S10: The concentrations of ribosomal proteins are more tightly regulated than non-ribosomal proteins**

A) The concentration increase in non-ribosomal proteins during shocks is correlated with the log of protein abundance during steady-state growth, unlike the ribosomal proteins.

B) The 40S ribosomal protein S2 is encoded by a single essential gene (*rps2*), and thus should be a better representation of the regulation of ribosome concentration. During more complete growth arrest with 1-M sorbitol shocks of 10-min period, its concentration increased only modestly (<20%). The dark centerline is the population-averaged cellular fluorescence (images acquired every 35 min), and the shading represents 1 S.D. ($n$ = 207 cells).

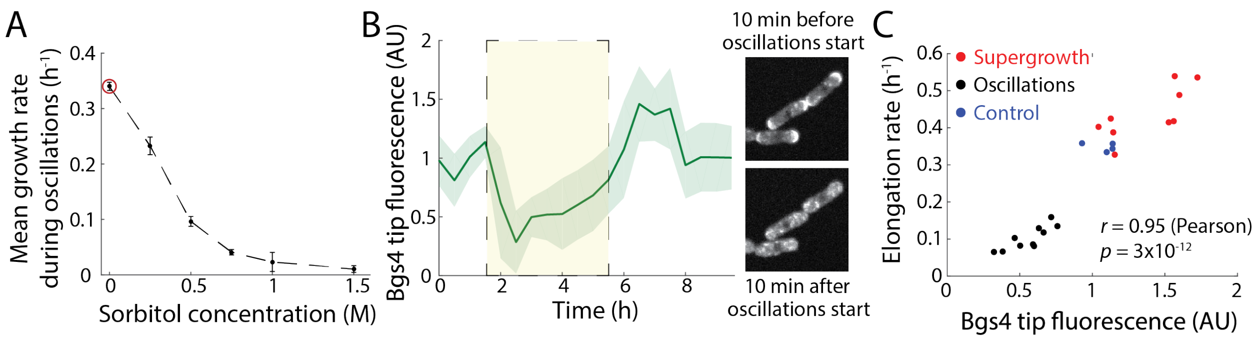


**Figure S11: Bgs4 tip concentration increases during supergrowth and correlates with elongation rate.**

A) Mean elongation rate during shocks was inversely related to the amplitude of osmotic shock with D-sorbitol during oscillatory cycles with 10-min period and 4-h duration. Error bars are 1 S.D. ($n$ = 265 cells).

B) Left: Localization of 2GFP-Bgs4 at the cell tips during an oscillatory osmotic shock experiment. Yellow outlined box represents the 4-h duration of 0.5 M sorbitol shocks with 10-min. Upon onset of oscillations, tip localization initially decreased by >70% (right), and re-localized concordant with the increase in growth rate during supergrowth ($n$ = 64 cells).

C) Population-averaged elongation rate was highly correlated with the level of 2GFP-Bgs4 tip localization during all phases of the experiment.

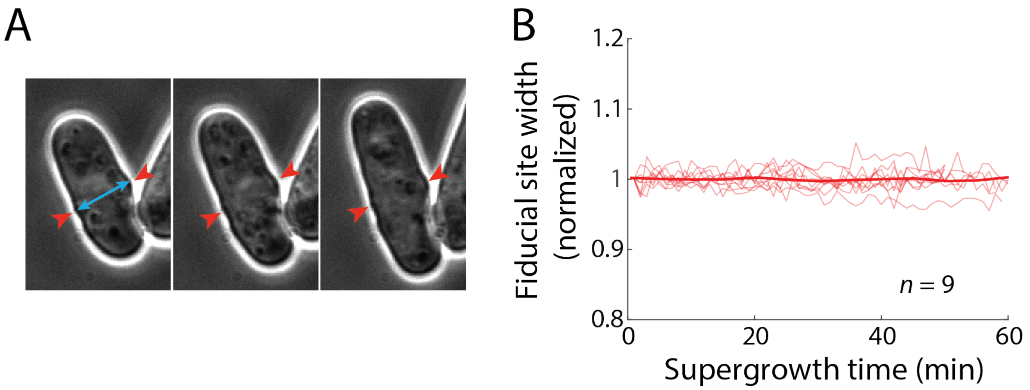


**Figure S12: Cells do not inflate or deflate during supergrowth**

A) Measurements of cell width at fiducial sites were used to assess changes in turgor. Red arrowheads mark a birth scar representing a previous cell division site, which remained identifiable throughout growth due to the lack of local cell-wall insertion.

B) Cell width measured from birth scars with high temporal resolution (10-s frame interval) remained approximately constant during supergrowth, indicating that turgor was not varying substantially ($n$ = 9). The dark red line indicates the average of all trajectories.

### Supplemental Movie Legends

**Movie S1: Wild-type *S. pombe* cells exhibit supergrowth after osmotic shock oscillations.** After 4 h of osmotic shock oscillations (0.5 M D-sorbitol, 10-min period, 24 cycles), cells exhibited supergrowth. Images were acquired with a frame interval of 1 min using phase-contrast imaging.

**Movie S2: The concentration of mCrimson increases during oscillations and dilutes during supergrowth.** FC3186 cells expressing mCrimson subjected to 4 h of osmotic shock oscillations (0.5 M D-sorbitol, 10-min period, 24 cycles), with subsequent supergrowth. Fluorescence images were acquired every 10 min (Methods).

**Movie S3: Treatment with the secretion-inhibiting drug brefeldin A results in increased mCrimson concentration and subsequent supergrowth.** FC3186 cells expressing mCrimson were subjected to treatment with 0.1 mg/mL brefeldin A for 4 h, during which time cell growth halted and mCrimson concentration increased. The drug was then washed out, and cells exhibited supergrowth and dilution of mCrimson.

### Supplementary Tables

#### Table S1: Strains used in this study.

| Strain | Genotype |
|---|---|
| FC15 | *h- WT (972)* |
| FC342 | *h- cdc25-22* |
| FC1037 | *h- sec6-GFP-ura4+ leu1-32 ura4-* |
| FC1503 | *h- ssp2::ura4+ leu1-32 ura4-D18* |
| FC1507 | *h- gad8::ura4+ leu1-32 ura4-D18* |
| FC1551 | *h- pmk1::ura4+ leu1-32 ura4-D18* |
| FC1568 | *h- sty1::ura4+ leu1-32 ura4-D18* |
| FC1596 | *h- cch1::ura4+ leu1-32 ura4-D18* |
| FC1897 | *h- fim1-meGFP-kanMX6* |
| FC1902 | *h- rga4-RFP::kanMX6 leu1-32 ura4-D18* |
| FC1919 | *h- ost1-GFP::ura4+ rlc1-RFP::ura4+ leu1-32 ura4-D18* |
| FC2086 | *h- trn1-GFP::kanMX6 ade6- leu1-32 ura4-D18* |
| FC2087 | *h- yop1-GFP::kanMX6 ade6- leu1-32 ura4-D18* |
| FC2255 | *h- leu1:2xGFP-bgs4 leu1+ ura4+* |
| FC2277 | *h- gma12-GFP-ura4+ leu1:tdTom-bgs1 leu1+ ura4+* |
| FC2678 | *h+ pom1-tomato-natMX, cdr2-GFP-kanMX ade6- leu1-32 ura4-D18* |
| FC2810 | *h- gpd1::kanMX leu1-32* |
| FC2840 | *h- tdh1-dendra2:ura4 ade6- leu1-32 ura4-D18* |

| | |
|---|---|
| FC2859 | *h+ hta1-mCherry:kanMX GFP-at2:kanMX ade6- leu1-32 ura4-* |
| FC2861 | *h+ GFP-atb2:kanMX ade6- leu1-32 ura4-D18* |
| FC2913 | *h- leu1-tomato-bgs1 psy1-GFP-leu1* |
| FC3186 | *h+ act1p::1XE2C:Hyg*$^{R}$ *ura4-D18 leu1-32 ade6-M210 his7-366* |
| FC3208 | *h- rps802-GFP::kanMX leu1-32 ura4-D18 ade6-M210* |
| FC3209 | *h- rps2-GFP::kanMX leu1-32 ura4-D18 ade6*$^{-}$ |
| FC3210 | *h+ rpl1601-GFP::kanR leu1-32 ura4-D18 ade6-M216* |
| FC3212 | *h- rpl2801-GFP::kanMX leu1-32 ura4-D18 ade6*$^{-}$ |
| FC3213 | *h+ rps2401-GFP::kanR leu1-32 ura4-D18 ade6-M216* |
| FC3215 | *h+ rpl3001-GFP::kanR leu1-32 ura4-D18 ade6-M216* |
